# Transite: A computational motif-based analysis platform that identifies RNA-binding proteins modulating changes in gene expression

**DOI:** 10.1101/416743

**Authors:** Konstantin Krismer, Shohreh Varmeh, Molly A. Bird, Anna Gattinger, Yi Wen Kong, Erika D. Handly, Thomas Bernwinkler, Daniel A. Anderson, Andreas Heinzel, Brian A. Joughin, Ian G. Cannell, Michael B. Yaffe

## Abstract

RNA-binding proteins (RBPs) play critical roles in regulating gene expression by modulating splicing, RNA stability, and protein translation. In response to various stimuli, alterations in RBP function contribute to global changes in gene expression, but identifying which specific RBPs are responsible for the observed changes in gene expression patterns remains an unmet need. Here, we present *Transite* a multi-pronged computational approach that systematically infers RBPs influencing gene expression changes through alterations in RNA stability and degradation. As a proof of principle, we applied Transite to public RNA expression data from human patients with non-small cell lung cancer whose tumors were sampled at diagnosis, or after recurrence following treatment with platinum-based chemotherapy. Transite implicated known RBP regulators of the DNA damage response and identified hnRNPC as a new modulator of chemotherapeutic resistance, which we subsequently validated experimentally. Transite serves as a generalizable framework for the identification of RBPs responsible for gene expression changes that drive cell-state transitions and adds additional value to the vast wealth of publicly-available gene expression data.

## I. Introduction

RNA-binding proteins (RBPs) are major modulators of gene expression at the post-transcriptional level, where they control RNA splicing, stability, localization, degradation, and translation [1,2]. RBPs play critical roles in cell differentiation and tissue development, and aberrant RBP function is implicated in a wide range of diseases, including neurodegenerative disorders and neuropathies, myopathies, autoimmune paraneoplastic syndromes, and cancer [3]. For mRNAs, the role of RBPs in modulating global changes in gene expression at both the RNA and protein level becomes particularly important under conditions where new gene transcription is repressed, such as during inflammation, cell stress, and in response to genomic damage [4–6]. Under these conditions, changes in gene expression have been shown to result, in part, from alterations in RBP activity [7]. Furthermore, mutations affecting the expression or function of specific RBPs have been implicated in a variety of diseases, including cancer [3,6,8,9].

RBPs recognize short linear sequence motifs containing 6 - 8 nucleotides within their target RNAs [10]. The identity of these motifs has been determined for a subset of all known RBPs using various *in vitro* based oligonucleotide selection methods such as SELEX [11], RNAcompete [12] and RNA Bind-n-Seq [13], and directly confirmed for a smaller set of RBPs through experimental analysis of RBP-RNA interactions using CLIP-seq and various extensions thereof. The RNA targets for most RBPs, as determined by CLIP-seq, however, have not been identified due to a variety of technical challenges, including cost, limited antibody specificity, and high background binding. Furthermore, direct experimental identification of RNA targets of RBPs likely depends on the experimental situation under which the CLIP-seq was performed. This lack of direct CLIP-seq data has limited our ability to directly map specific RBPs onto global changes in RNA levels, including those in patient-based gene expression data sets, that have been observed following various stimuli or clinical treatments.

RBPs appear to play a particularly important role in orchestrating the DNA damage response (DDR) by regulating mRNA expression changes that control the onset and duration of cell cycle checkpoints and drive DNA repair [14–16]. Unbiased large-scale screening efforts have converged on RBPs as one of the most enriched classes of proteins modulating the DDR, even more so than annotated DNA damage repair proteins [17–21]. In addition, emerging evidence from a number of labs has identified RBPs as critical targets of DDR kinases, including both upstream sensor kinases such as ATM, ATR and DNA-PK, and downstream effector kinases such as Chk1 and MK2 [17,18,22–24]. The discovery of RBPs as integration points of the cellular response to genomic damage has important clinical applications, since the efficacy of many commonly used chemotherapeutic drugs is dependent on the integrity (or lack thereof) of the DDR [25,26]. For example, we found that a key target of the DNA damage-activated MK2 pathway was the RBP hnRNPA0, which was required for maintenance of the G1/S and G2/M checkpoints following cisplatin treatment [27,28]. Furthermore, this finding has clear clinical relevance to the response of non-small cell lung cancers (NSCLCs) to chemotherapy in both mouse models and human patients, where the expression levels of two critical hnRNPA0 target RNAs, Gadd45α and p27, predicted the clinical response of mouse and human tumors to platinum therapy. Despite these types of data, and the recent surge of interest in the roles of RBPs in cancer chemosensitivity and resistance [6,14,29], general methods for the systematic identification and prioritization of RBPs that influence various biological responses, including the DDR in clinically relevant patient-based gene expression data sets, are lacking.

To address this we developed a computational approach, called *Transite*, that leverages pre-existing gene expression data and known RBP binding preferences in order to infer RBPs that may be responsible for alterations in RNA levels under a given condition or perturbation. This approach is analogous to our previous computational tool *Scansite*, which predicts substrates of kinases and modular signaling domains based on phosphorylation and peptide-binding motifs [30]. With Transite, we hope to expand the utility of RBP biology to the wider scientific community.

## II. Results

The overall approach used by Transite to map RBPs to sets of differentially expressed genes is illustrated in Figure 1. Transite starts with a list of differentially expressed genes between two conditions (i.e. treated versus untreated samples), identifies short linear oligonucleotide motifs or *k*-mers that are enriched or depleted within specific regions of the transcripts they encode (i.e. 5′-UTR, CDS, or 3′-UTR), and then matches these motifs to likely RBPs that bind them using a compendium of known RBP motifs (see IV-B). Transite’s default setting is to analyze 3′-UTR sequences, since motifs that regulate mRNA stability typically reside within the 3′-UTR, but also allows the same analysis to be performed on the CDS or the 5′-UTR. Two different approaches are used, depending on whether the set of differentially expressed genes is first separated into distinct foreground and background sets, or instead is analyzed as a continuous list of genes ordered by change in expression level. For the former approach in which foreground sets are pre-determined by differentially expressed genes, we developed Transcript Set Motif Analysis (TSMA), which looks for enriched or depleted oligonucleotide motifs based on systematic differences between the foreground sets and the total gene expression data (i.e. the background). For the latter approach (i.e. a list of ranked genes) we developed Spectrum Motif Analysis (SPMA), which analyzes motif enrichment along that ordered list of transcripts, similar to the approach taken by Gene Set Enrichment Analysis [31]. This approach exploits information across the entire spectrum of changes rather than limiting analysis to the up- and downregulated extremes, and allows motif enrichment or depletion to be visually displayed as a color spectrum. Both TSMA and SPMA then use two distinct methods, a *k*-mer-based and a matrix-based method, to score for and infer candidate RBP in the differentially expressed genes. The *k*-mer-based and matrix-based implementations of TSMA and SPMA are explained in more detail below.

**Fig. 1.**
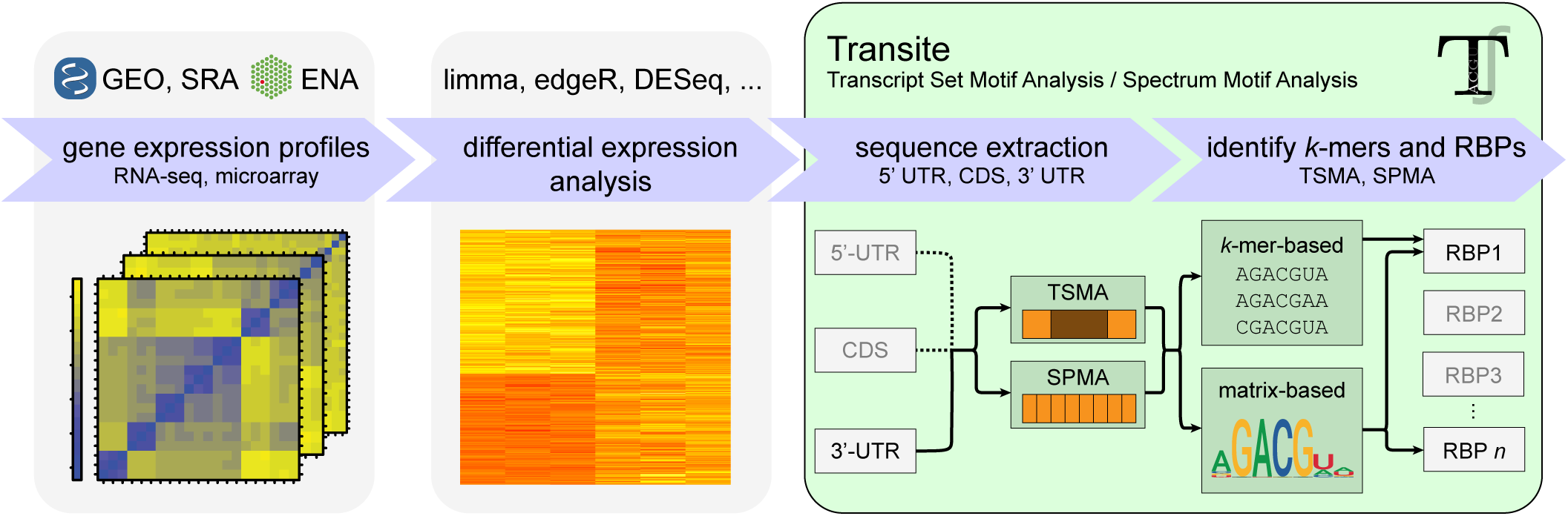
Schematic figure of the Transite analysis pipeline. The initial steps of the Transite data analysis workflow include preprocessing and differential expression analysis of gene expression profiles, which could be collected in-house or obtained from NCBI and EMBL-EBI repositories such as GEO, SRA, and ENA. Differential expression analysis is used to either identify groups of upregulated and downregulated genes (for Transcript Set Motif Analysis) or to establish a ranked list of genes from most upregulated to most downregulated (for Spectrum Motif Analysis). Transite then analyzes regions within these genes to identify *k*-mers and RBPs whose motifs are enriched or depleted in the differentially expressed genes.

### A. *Transcript Set Motif Analysis identifies enriched and depleted* k*-mers within assigned sets of upregulated and downregulated genes and maps them onto RBPs*

Transcript Set Motif Analysis identifies the overrepresentation or underrepresentation of all possible hexamers or heptamers, as well as binding motifs for 174 well-characterized RNA-binding proteins in a set (or sets) of transcripts (i.e. a foreground set), relative to the background of the entire population of transcripts measured in an experiment (Figure 2A).

**Fig. 2.**
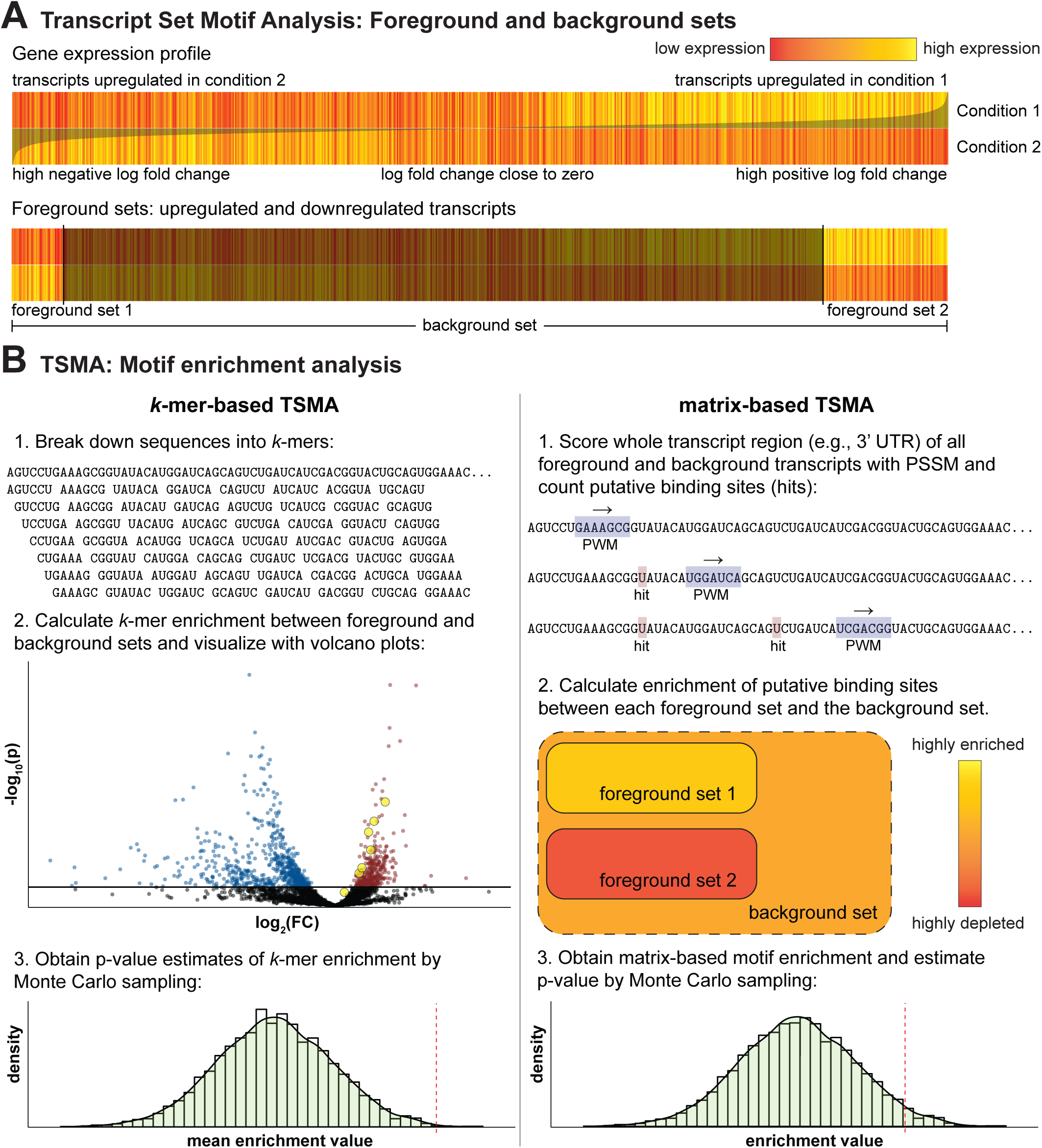
Transcript Set Motif Analysis. (**A**) Foreground sets in TSMA are defined by differential gene expression analysis of RNA-seq or microarray data sets, usually by selecting statistically significantly upregulated and downregulated genes. The background set is all genes in the microarray platform or all measured genes in RNA-seq. In the heatmap of the gene expression profile in panel **A**, the two rows (*Condition 1, Condition 2*) are the mean gene expression values of the replicates of the respective groups (e.g., *Condition 1* could be *treated with drug A* and *Condition 2 untreated*). The columns of the heatmap correspond to the genes, and the superimposed gray curve is the log fold change between *Condition 1* and *Condition 2*. (**B**) TSMA estimates the statistical significance of putative RBP binding site enrichment between each foreground set and the background set. There are two ways to describe putative binding sites of RNA-binding proteins (i.e. the motif). The column on the left depicts *k*-mer-based TSMA, which uses a list of *k*-mers to describe putative binding sites. The column on the right is matrix-based TSMA, which instead uses Position Weight Matrices (PWMs). See text for details.

Two different methods are used to assign transcript targets to specific RBPs. One of the methods, *k*-mer-based TSMA, also identifies statistically significantly overrepresented and underrepresented hexamers or heptamers within the fore-ground set, irrespective of whether they can be associated with a known RBP motif. Matrix-based TSMA leverages the full PWM representations (see IV-C for details) of known RBP motifs to nominate RBPs whose motifs are overrepresented or underrepresented in the foreground set.

In the *k*-mer-based approach, after foreground and background sets are defined (Figure 2A) and the preferred sequence region is selected (5′-UTR, CDS, or 3′-UTR), the sequences of both sets are broken down into overlapping hexamers or heptamers (i.e. *k*-mers of length 6 or 7, respectively) (Figure 2B, left column, step 1), and for each *k*-mer its frequencies in the foreground and background set are determined. While Transite supports both hexamer- and heptamer-matching, hexamers are recommended, since computer run-time increases exponentially with *k* and the results for heptamers mirror those for hexamers in our experience.

The enrichment value of a particular *k*-mer *i, e*_*i*_, is then calculated as follows:

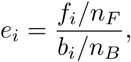

where *f*_*i*_ and *b*_*i*_ are the absolute counts of *k*-mer *i* in foreground and background set and *n*_*F*_ and *n*_*B*_ are the total counts of all *k*-mers in the foreground and background, respectively.

The statistical significance of the enrichment for each *k*-mer is then determined. First, a contingency table *C*_*i*_ for *k*-mer *i* is defined as

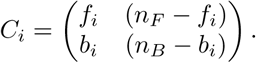

Then, the p-value *p*_*i*_ for *C*_*i*_ is approximated with Pearson’s *χ*^2^ test. If *p*_*i*_ *<* 5*α*, where *α* is the decision boundary, and *p*_*i*_ is replaced by the p-value obtained by Fisher’s exact test for *C*_*i*_. This step-wise procedure reduces computation time dramatically (approximately 50-fold), because the computationally expensive Fisher’s exact test is only used in cases where the approximate p-value from the computationally inexpensive *χ*^2^ test is close to the decision boundary and is avoided in cases where a precise p-value is unnecessary. Furthermore, Fisher’s exact test is always used if at least one of the expected counts is less than five, because this constitutes a violation of the assumptions of the approximate test. The p-values are subsequently adjusted for multiple hypothesis testing. The available p-value adjustment methods are described in section 5 of Supplementary Methods.

The list of all *k*-mers with their associated enrichment values and statistical significance in the foreground sets is then reported. This is particularly important because it provides an unbiased way to identify overrepresented and underrepresented sequences and novel motifs regardless of whether they conform to known RBP binding motifs. The results are visualized using volcano plots that show the enrichment values on the x-coordinate (log transformed) and the associated p-values on the y-coordinate (log transformed and multiplied by −1) for all *k*-mers. An example is shown in Figure 2B, where the black dots represent *k*-mers without significant enrichment or depletion, while blue dots denote significantly depleted *k*-mers and red dots significantly enriched *k*-mers. The *k*-mers corresponding to the motif of one particular RBP are indicated by yellow circles.

Over- and under-represented *k*-mers are then mapped onto specific RBPs. A set of *k*-mers associated of each RBP is generated from the known RBP motif PWMs, as described in IV-C. These RBP-specific *k*-mers are then assigned the enrichment values calculated from the data, as shown by the yellow dots in the volcano plot in Figure 2B. The geometric mean of the enrichment values of all *k*-mers that are associated with that particular RBP is then calculated, and analyzed for its statistical significance using Monte Carlo sampling (see section 2 in Supplementary Methods). A null distribution of mean enrichment values associated with an RBP’s *k*-mers is generated by repeated random selection of foreground sets from the background. The null distribution is used to obtain an estimate of the significance of the true mean enrichment value of the RBP-associated *k*-mers observed in the experimental data, which is shown as a red dashed line in the histogram in Figure 2B, step 3. A ranked list of RBPs and their associated p-values, corrected for multiple hypothesis testing, is then provided.

An alternative to *k*-mer-based TSMA is a matrix-based approach, where the sequence motifs of 174 RBPs are maintained as PWMs. All sequence positions in the transcripts within the foreground and background gene sets are then scored, as shown in step 1 of the right column of Figure 2B. The PWM slides along the sequence, assigns a score to each position, and scores above a certain threshold are considered putative binding sites (*hits*), (see section 1 in Supplementary Methods). These hits are tallied in both the foreground and the background set, and enrichment values and associated p-values calculated analogously to the *k*-mer-based approach. Again, all p-values are multiple testing corrected.

One disadvantage of the matrix-based TSMA method relative to the *k*-mer-based approach is that a PWM assumes independence among positions, making it impossible to construct a PWM that assigns high scores to AAAAAA and CCCCCC, but a low score to ACACAC. An advantage of our matrix-based approach, however, is it retains positional hit information within the sequence and therefore facilitates the detection of closely spaced clusters of putative binding sites. Homotypic clusters of binding sites on DNA, for example, have been shown to be important for transcription factor binding [32], and have been postulated to be involved in RNA regulation [33,34], but a clear experimental demonstration of their general importance for RBP binding to RNA has not been unambiguously shown.

### B. Spectrum Motif Analysis identifies RBPs with non-random arrangement of putative binding sites in a ranked list of transcripts

A limitation of the TSMA method described above is that it will only capture those RBPs for which putative binding sites are statistically significantly enriched among a pre-defined foreground set of differentially expressed genes relevant to a background set. As an alternative method, we developed Spectrum Motif Analysis (SPMA), an approach that more broadly and generally identifies non-random distributions of RBP target sites in an ordered list of genes without having to pre-define a specific foreground set (compare Figures 2A and 3A).

**Fig. 3.**
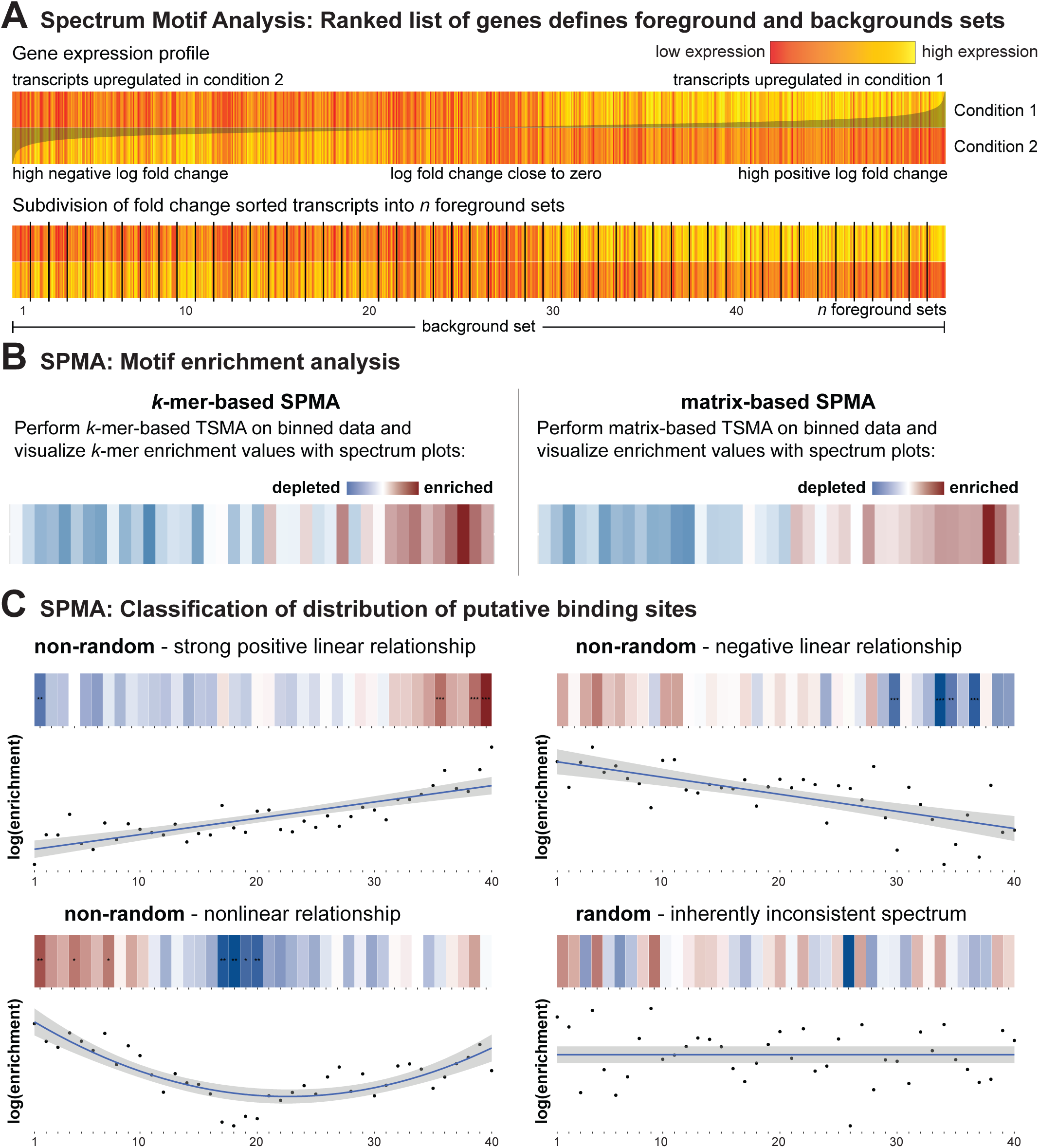
Spectrum Motif Analysis. (**A**) Transcripts are sorted by some measure of differential expression (e.g., fold change or signal-to-noise ratio) and the entire spectrum of transcripts is then subdivided into a number of equally-sized foreground “bins”. (**B**) The motif enrichment step is identical to TSMA. SPMA results are visualized as spectrum plots, which are one-dimensional heatmaps of motif enrichment values, where the columns correspond to the bins and the color encodes the enrichment value (strong depletion in dark blue to strong enrichment in dark red) of a particular *k*-mer or PWM. (**C**) The distribution of putative binding sites (as visualized by spectrum plots) is deemed *random* or *non-random* (i.e. putative binding sites are distributed in a way that suggest biological relevance), based on multiple criteria described in the text. Shown beneath each strip in the heat map are the log enrichment values for the RBP motif being analyzed (black dots), and the best first, second, or zero order polynomial fit (blue line) along with 95% confidence intervals (shaded gray).

Instead of using an arbitrary threshold (e.g., p-value less than or equal to 0.05) to assign transcripts to a single foreground set, SPMA subdivides the entire list of rank-ordered transcripts into a number of bins of equal width. Each bin is considered its own foreground set and enrichment scores for *k*-mers or PWM motifs are then calculated as described above. The enrichment scores for each RBP across the bins are then visualized as one-dimensional heatmaps, where red-blue coloring encodes the putative binding site enrichment values, as shown in Figure 3B, to generate spectrum plots. RBPs that are involved in regulating differential gene expression should show non-random red-blue color patterns in the spectrum plot, indicating progressive RBP binding motif enrichment in the upregulated genes, the downregulated genes, or both. As shown in the upper left plot of Figure 3C, genes that are upregulated in condition 1 show a progressive overrepresentation of putative binding sites for a particular RBP, consistent with that RBP enhancing mRNA stability. In contrast, as shown in the upper right plot of the same panel, genes that are downregulated in condition 1 show a progressive overrepresentation of binding sites for a different RBP, consistent with this RBP destabilizing its mRNA targets.

SPMA generates one spectrum plot for each RBP motif in the motif database. With 174 motifs currently available, it is imperative to provide an analytical means to aid in the identification of biologically meaningful spectrum plots that exhibit non-random patterns. Each spectrum plot is therefore examined for whether the distribution of enrichment values among the bins is *non-random* or *random*, based on three criteria: (1) the adjusted *R*^2^ of a polynomial model fit, (2) the local consistency score, and (3) the number of bins with a significant enrichment or depletion of putative binding sites. The significance of the enrichment values is calculated in an identical fashion to the significance calculation in TSMA. For (1), polynomial regression models of degrees ranging from 0 through 5 are fitted to the spectrum of enrichment values, and the model that best reflects the true nature of the data is selected by means of the F-test (see section 6.2 of Supplementary Methods for details on the polynomial model approach). Examining the coefficient of the linear term in the polynomial depicts the general increase or decrease in RBP enrichment along the bins as illustrated in the first two examples of Figure 3C, respectively. If there is a strong evidence for a non-linear relationship, this can also be captured by the model, as seen in the third example shown in Figure 3C. With approach (2), a local consistency score quantifies the local noise of the spectrum by calculating the deviance between the linear interpolation of the scores of two bins separated by exactly one other, and the observed score of the middle bin, for each position in the spectrum. The lower the score, the more consistent the trend in the spectrum plot (see section 6.1 of Supplementary Methods for a formal definition of the local consistency score and section 2 for details on the Monte Carlo sampling procedure of the null distribution of the score). Spectrum plots are classified as non-random if (1) the adjusted *R*^2^ of the polynomial fit is greater than or equal to 0.4, and (2) the p-value associated with the local consistency score is less than or equal to 5 ∗ 10^−6^, and (3) at least 10% of the bins have significant (*α* = 0.05) enrichment or depletion of putative binding sites.

### C. Website and R package for Transite available for customizable use

To make RBP analysis of gene expression data sets widely available to the scientific community, the Transite analysis platform is hosted at https://transite.mit.edu. Both the TSMA and SPMA methods are web-accessible and familiarity with the R programming language is not required (Figure 4). The full functionality of Transite is also provided as an R/Bioconductor package (https://doi.org/10.18129/B9.bioc.transite) to facilitate a seamless integration of these algorithms into existing bioinformatics workflows. The source code of the Transite package is hosted on GitHub (https://github.com/kkrismer/transite). Both website and the R package also allow motif enrichment analysis with user-defined motifs, in addition to the 174 motifs provided by the Transite motif database, enabling users to search for enrichment of any RBP motif in a discrete set of genes or a rank-ordered list.

**Fig. 4.**
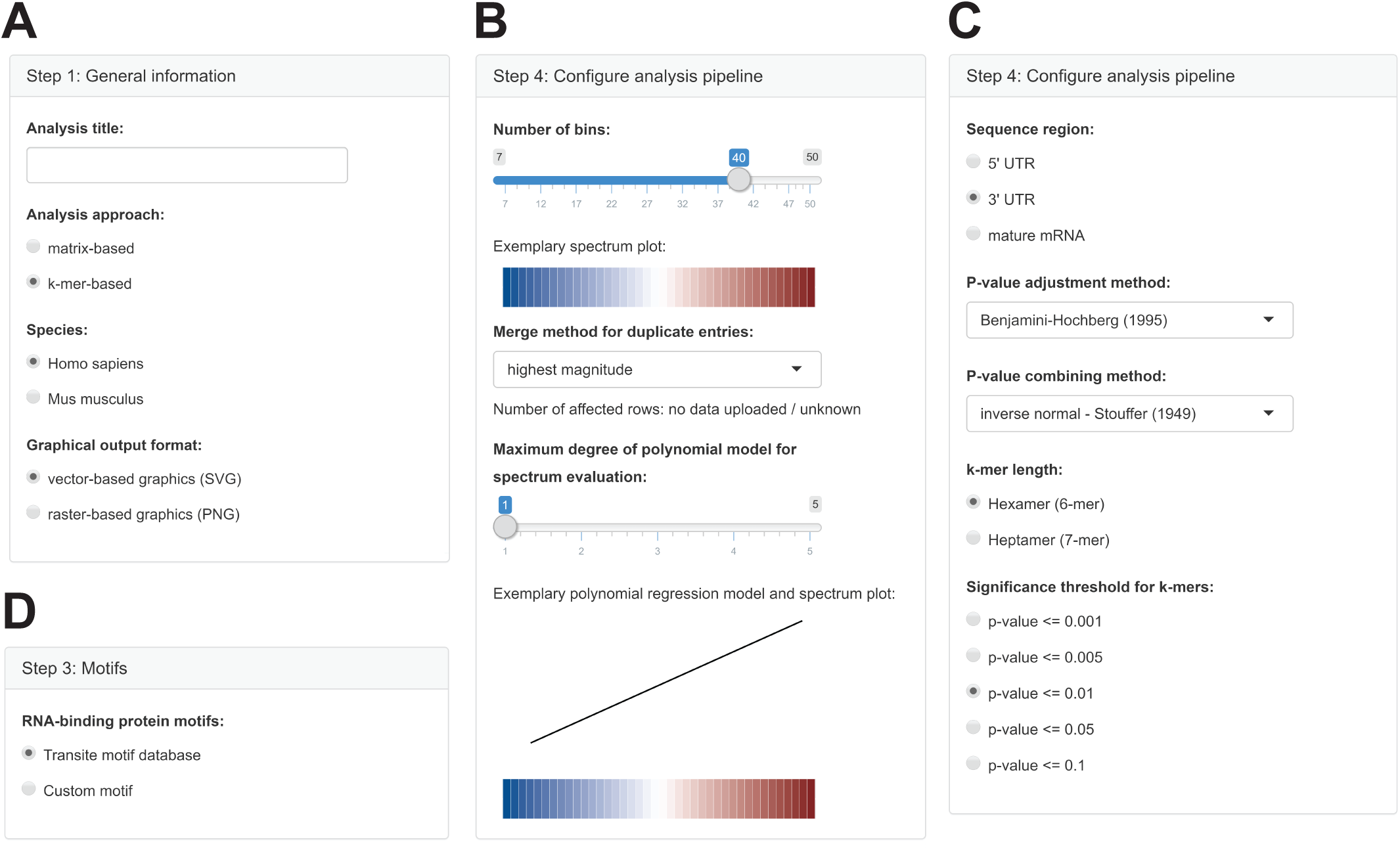
Transite web interface. Data sets are analyzed using TSMA or SPMA in four simple steps, some of which are illustrated in panels **A** - **D**. These involve the selection of *k*-mer or matrix-based analysis (**A**), the specification of foreground and background sets for TSMA, the number of bins for SPMA (**B**), the region of the RNA to be analyzed and the threshold for statistical significance (**C**), and the source of RNA binding motifs to be used for the analysis (**D**).

### D. Transite correctly maps observed changes in RNA abundance following ZFP36 overexpression or ELAVL1 knockdown onto their respective RBPs

To test the ability of the Transite algorithms to correctly map changes in RNA expression onto specific RBPs, we used a publicly available data set in which RNA expression levels were measured following overexpression of the RBP ZFP36 (also known as TTP). ZFP36 is known to destabilize its target RNA transcripts by binding to sequence elements in the 3′-UTR [35]. Mukherjee et al. (2014) reported microarray measurements of differential RNA expression in HEK293 cells following inducible overexpression of an EGFP-ZFP36 fusion protein (GEO series accession GSE53185). The RNA expression fold change and associated p-values per gene between the induced and un-induced groups, as reported by the authors, were used as input for Transite. Genes that were statistically significantly downregulated and upregulated following ZFP36 overexpression (i.e. *p <* 0.05 after multiple testing correction) were chosen as foreground sets for TSMA. Volcano plots showing *k*-mer enrichment and depletion in these gene sets are shown in Figure 5A, and the top 10 empirically identified *k*-mers are listed in Supplementary Tables S1 and S2. The left panel in Figure 5A shows that *k*-mers corresponding to the ZFP36 binding motif, shown in yellow, are among the most highly enriched *k*-mers in transcripts that were found to be downregulated, while the right panel shows conversely that ZFP36 associated *k*-mers were highly depleted in the genes that were upregulated after ZFP36 overexpression. This was even more apparent in the spectrum plot following SPMA of this data set (Figure 5B), which revealed a highly consistent nearly monotonic increase in ZFP36 binding sites when the genes were ranked from those most upregulated to those most downregulated after ZFP36 overexpression. On this basis, ZFP36 emerged as the single best RBP out of all 174 RBPs in the database whose motif could rationalize the observed gene expression changes.

**Fig. 5.**
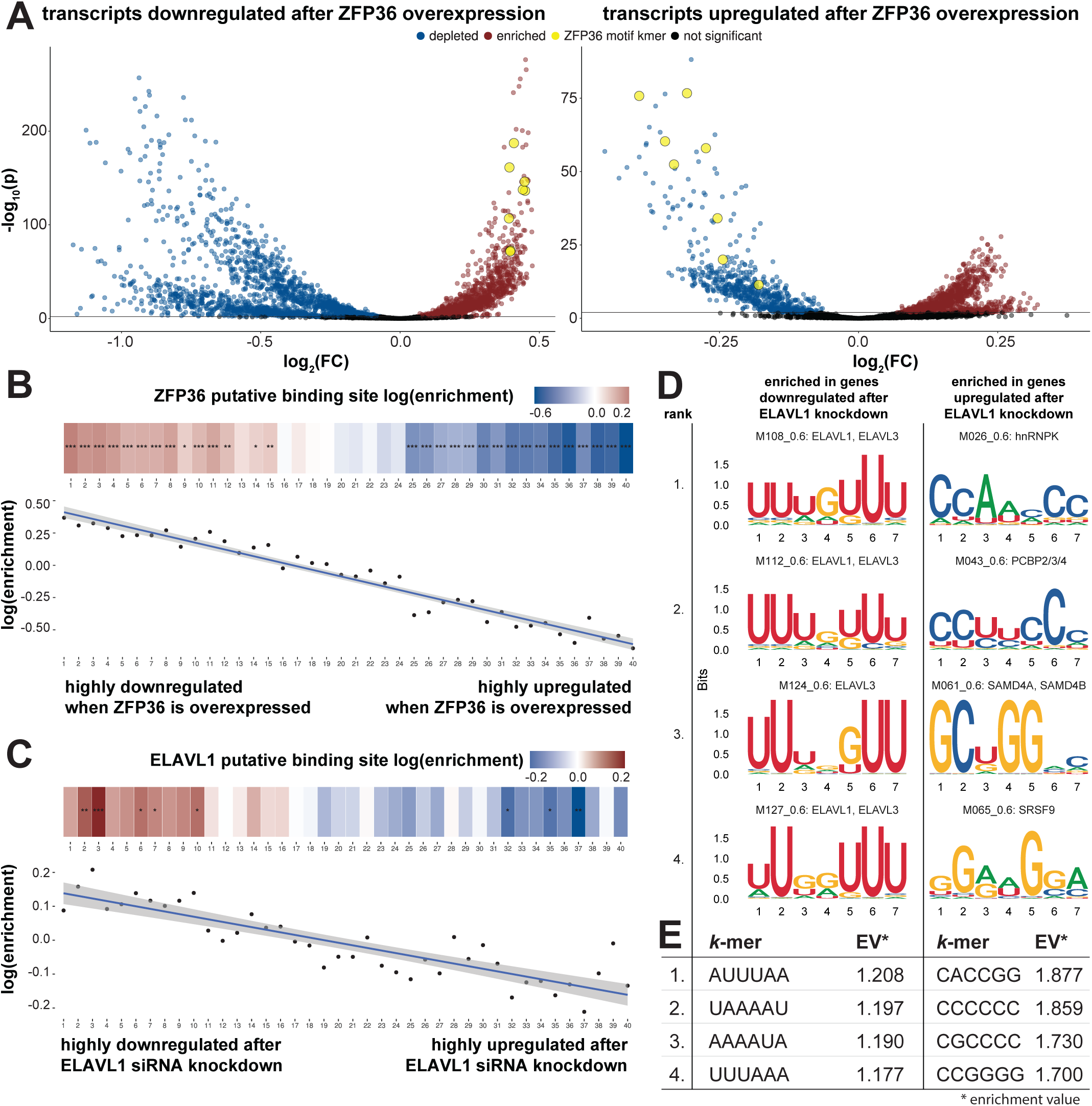
Unbiased identification of drivers of differential expression after overexpression of ZFP36 or knockdown of ELAVL1. (**A**) TSMA volcano plot showing enriched and depleted *k*-mers in downregulated transcripts after ZFP36 overexpression (right panel). *k*-mers associated with ZFP36 (shown in yellow) are highly enriched. TSMA volcano plot of *k*-mer enrichment values in upregulated transcripts after ZFP36 overexpression shows strong depletion of ZFP36 associated *k*-mers (right panel). (**B**) SPMA spectrum plot depicts relationship between ZFP36 overexpression and downregulation of ZFP36 targets. (**C**) SPMA spectrum plot of one ELAVL1 motif depicting global downregulation of ELAVL1 target transcripts after ELAVL1 siRNA knockdown. (**D**) Sequence logos of motifs highly enriched in transcripts upregulated (left column) and downregulated (right column) after ELAVL1 knockdown. U-rich ELAVL1 motifs are highly enriched in the 3′-UTRs of downregulated transcripts (GSE29778). (**E**) Four most highly enriched hexamers in transcripts upregulated (left column) and downregulated (right column) afer ELAVL1 knockdown, as identified by *k*-mer-based TSMA.

To further validate the utility of Transite to infer RBPs that modulate gene expression changes, we used a second publicly available data set (GEO series accession GSE29778) in which gene expression changes were measured following siRNA knockdown of ELAVL1 (also known as HuR) to 20% of its endogenous levels [34]. ELAVL1 stabilizes its target RNA transcripts and likely facilitates their pre-mRNA processing, hence its knockdown should result in reduced expression of its target RNAs. As shown in Figure 5C, analysis of this data set using SPMA resulted in spectrum plots in which the enrichment values for the ELAVL1 motifs closely varied in direct proportion to the extent of RNA downregulation that was observed (Figure 5C and Supplementary Figure S1). Figure 5D shows the top 5 RBP motifs that were enriched in the upregulated and downregulated genes, revealing that genes downregulated after ELAVL1 knockdown were enriched in U-rich RBP motifs, including those that correspond to the ELAVL1 motifs in the Transite motif database. In contrast, genes that were upregulated after ELAVL1 knockdown were enriched in alternative RBP motifs that lacked U-rich regions, and corresponded to the binding motifs of other RBPs. Furthermore, the single most highly enriched *k*-mer in the set of downregulated genes, AUUUAA, that was empirically identified by *k*-mer-based TSMA (Figure 5E and Supplementary Tables S3 and S4), perfectly matches the motif of ELAVL1 that was experimentally determined using PAR-CLIP and RIP-chip [34]. Taken together, these data indicate that Transite can capture the specific RBPs responsible for gene expression changes caused by manipulation of RBP levels, thus validating our approach and providing confidence that predictions derived from more complex perturbations are more likely to reflect real changes in RBP binding or activity.

### E. RBPs involved in the DNA damage response are identified by Transite using cancer patient RNA expression data

As an application of Transite-based RBP scoring, we next analyzed a gene expression data set from patients with non-small cell lung cancer (NSCLC), who were either treatment-naive, or had recurred after platinum-based chemotherapy treatment (GEO series accession GSE7880). Differences in RNA transcript abundance were ranked between the set of tumors that were sampled pre-treatment and the separate set of tumors that were sampled after recurrence following treatment, and the ranked transcripts then analyzed by Transite in order to identify potential RBPs that might influence the response to platinum treatment. Changes in transcript abundance were ranked based on signal-to-noise ratio where transcripts upregulated in recurrent patients had positive values and those upregulated in treatment-naive patients had negative values (see Figure 6A for schematic). *k*-mer-based TSMA, focusing on the 3′-UTRs of the differentially regulated genes, revealed a set of enriched *k*-mers in the patients whose tumors failed platinum treatment that were largely U-rich (Supplementary Table S5). These *k*-mers mapped to the motifs of ELAVL1 and TIA1 as the top 2 hits (Figure 6B, top). SPMA revealed these same top two RBPs, as shown in the bottom part of Figure 6B. Individual spectrum plots for ELAVL1 (Figure 6C) and TIA1 (Figure 6D) demonstrated consistent behavior of these motifs across the gene expression continuum, being enriched in 3′-UTRs of genes that were upregulated in patients with recurrent tumors after platinum treatment, and depleted in 3′-UTRs of genes that were upregulated in naive patients. Importantly, upregulation of ELAVL1 and TIA1-target mRNAs was further validated by analyzing the distribution of previously known CLIP-Seq identified targets [36,37] for these two RBPs (Figure 6E and 6F), suggesting that our motif-based approach can identify bona fide target genes of a given RBP for which CLIP-Seq data is available. Moreover, both ELAVL1 and TIA1 are known to be involved in the DNA damage response [38–41]. The fact that two well-known players in the DNA damage response were among the top hits of the motif analysis provides confidence that Transite’s predictions are likely to reflect regulators of the DNA damage response and drivers of chemoresistance.

**Fig. 6.**
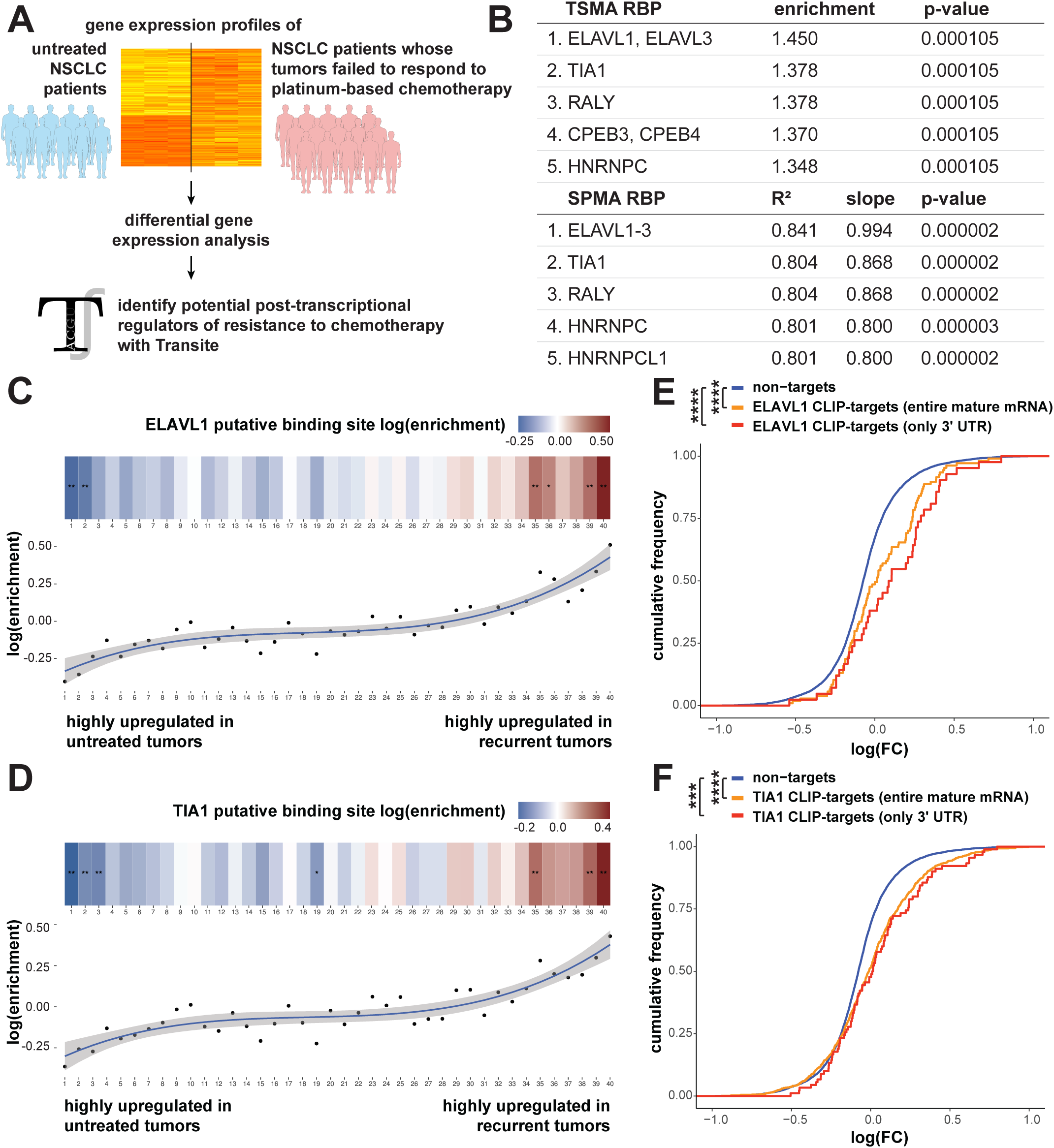
SPMA identifies ELAVL1 and TIA1 motifs as highly enriched in recurrent NSCLC patients. (**A**) Differential gene expression analysis was performed on samples from patients with untreated NSCLC tumors and patients with recurrent tumors. (**B**) Transite was used to identify RBPs whose targets were overrepresented among upregulated genes in samples of recurrent tumors. Shown are two tables of *k*-mer-based TSMA and SPMA showing RBPs with highly enriched motifs for TSMA and highly non-random motif enrichment pattern for SPMA. Among the top hits are ELAVL1, TIA1, and hnRNPC. (**C**) Spectrum plot from SPMA depicting the distribution of putative ELAVL1 binding sites across all transcripts. The transcripts are sorted by ascending signal-to-noise ratio. Transcripts downregulated in resistant samples relative to untreated samples are on the left, and those upregulated are on the right of the spectrum. Putative binding sites of ELAVL1 are highly enriched in transcripts upregulated in resistant cells (shown in red) and highly depleted in transcripts downregulated in resistant cells (shown in blue). (**D**) Spectrum plot of putative TIA1 binding sites using same transcript order as in panel C. (**E**) Enrichment of ELAVL1 targets in resistant NSCLC cells is recapitulated in an independent HITS-CLIP experiment (publicly available data). The distribution of fold changes of transcripts that have ELAVL1 binding sites is shifted in the positive direction, even more so when the binding sites are in the 3′-UTR. The p-values were calculated with the one-sided Kolmogorov-Smirnov test. (**F**) As in panel E, transcripts with TIA1 binding sites are upregulated in resistant cells according to an iCLIP experimenmt, confirming results from SPMA.

### F. Motif analysis of recurrent non-small cell lung cancers after cisplatin treatment identifies hnRNPC as a potential modulator of drug resistance

We were particularly interested in using Transite as a tool to identify new RBPs potentially involved in chemosensitivity or resistance to DNA-damaging chemotherapy agents using data from human clinical trials. We therefore chose to focus on hnRNPC, one of the highest-scoring RBPs that emerged from both TSMA and SPMA analysis of chemoresistant NSCLC patients, and one that has not, to our knowledge, been strongly implicated in the response to chemotherapy-induced DNA damage [42]. As shown in Figure 7A, the spectrum plot of the distribution of putative hnRNPC binding sites shows a strong enrichment of mRNAs with hnRNPC motifs in their 3′-UTRs in patients whose tumors recurred after platinum therapy. This Transite prediction was independently confirmed by analysis of iCLIP-defined target mRNAs for hnRNPC [43], which also showed an overrepresentation of hnRNPC targets in upregulated transcripts in recurrent patients (Figure 7B), with those with binding in the 3′-UTR showing the strongest enrichment.

**Fig. 7.**
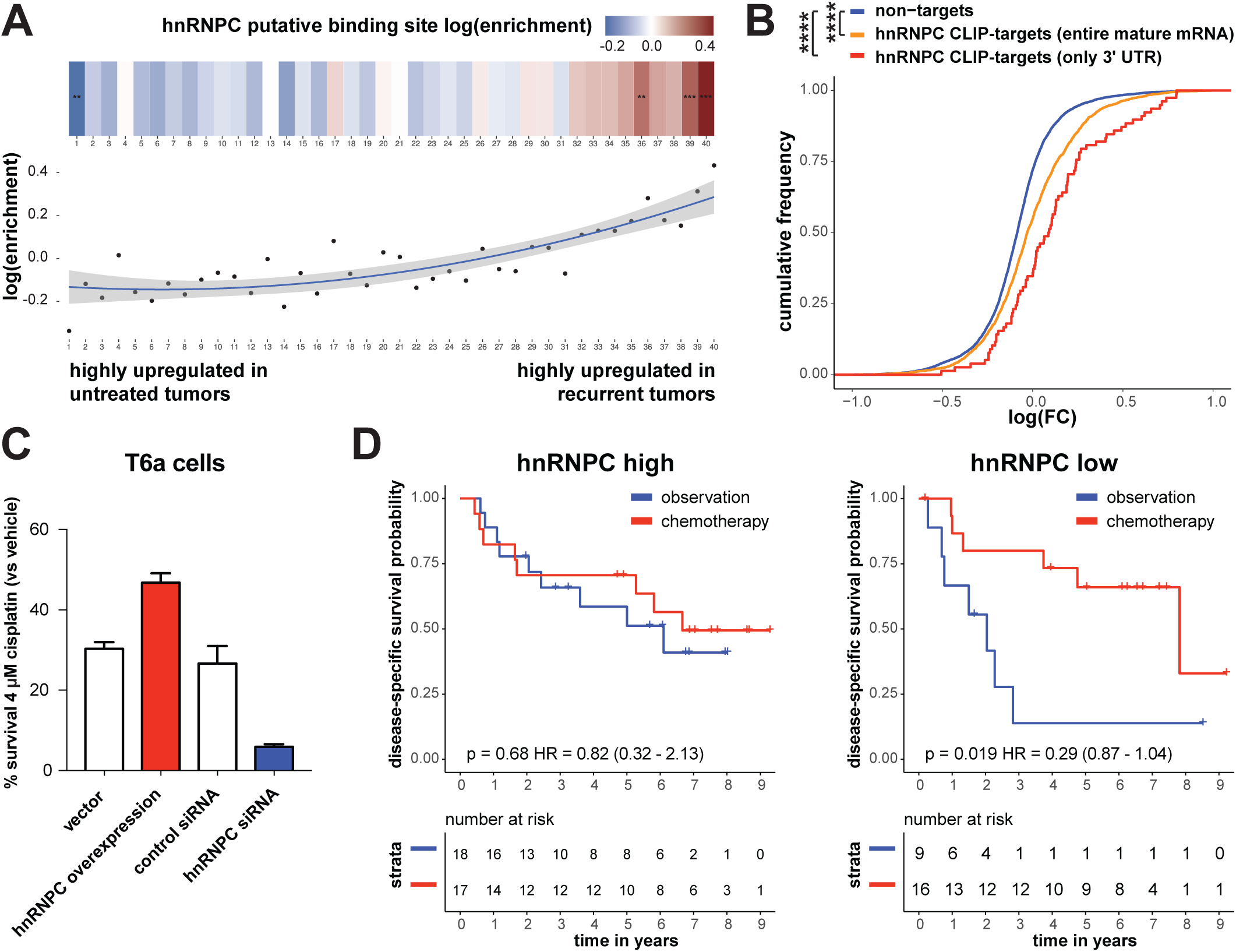
hnRNPC modulates sensitivity to cisplatin. (**A**) Spectrum plot from *k*-mer-based SPMA depicting the distribution of putative hnRNPC binding sites across all transcripts in samples from patients with untreated NSCLC tumors and patients with recurrent tumors as in Figure 6. The transcripts are sorted by ascending signal-to-noise ratio from lowest to highest abundance in resistant relative to untreated samples. Putative hnRNPC binding sites are highly enriched in the upregulated fraction of transcripts (GSE7880). (**B**) Enrichment of hnRNPC binding sites in upregulated transcripts is independently confirmed by CLIP experiments. The p-values were calculated with the one-sided Kolmogorov-Smirnov test. (**C**) siRNA-mediated reduction in hnRNPC levels significantly impairs long-term survival of T6a cells in response to cisplatin (blue bar). Overexpression of hnRNPC (red bar) protects against cisplatin-induced cell death in T6a cells in colony formation assays. Bar graphs represent percent number of colonies formed, normalized to untreated control cells. White bars represent control cells transfected with control vehicles (control siRNA or empty pcDNA). Error bars represent standard deviation among 3 replicates. (**D**) High expression of hnRNPC are associated with decreased efficacy of platinum-based chemotherapy in patients with stage 2 disease from the JBR.10 lung cancer adjuvant chemotherapy trial (GSE14814). The p-value was calculated with the log-rank test (HR is Hazard Ratio). hnRNPC low group = patients with hnRNPC expression Z-scores of less than or equal to −0.2, and hnRNPC high group = patients with hnRNPC expression Z-scores greater than or equal to 0.2.

To experimentally test these Transite predictions, we examined the effect of knockdown or overexpression of hn-RNPC in T6a murine lung carcinoma cells on their sensitivity and resistance to cisplatin treatment. As shown in Figure 7C, colony formation assays in T6a cells demonstrated that hnRNPC overexpression promoted resistance to cisplatin as evidenced by a 1.6 fold increase in the number of surviving colonies (Figure 7C, red bar). Conversely, siRNA-downregulation of hnRNPC significantly enhanced T6a cell sensitivity to cisplatin as evidenced by a 5-fold decrease in the number of colonies formed by cells treated with hnRNPC siRNA compared to those of control siRNA-treated cells after cisplatin treatment (Figure 7C, blue bar). These data indicate that hnRNPC mediates resistance of NSCLC cells to cisplatin chemotherapy, consistent with what was seen in the patient data, and demonstrate that our computational approach can identify new RBPs influencing the DDR.

To independently validate the importance of hnRNPC in mediating chemotherapy response in patients, we took advantage of data from a unique adjuvant chemotherapy trial, JBR.10 (Figure 7D) [44]. In this trial, early stage NSCLC patients had their tumors surgically resected and subjected to gene expression profiling (GEO series accession GSE14814). Patients were then randomized to receive cisplatin/vinorelbine combination chemotherapy or observation and palliative care, allowing us to specifically query the role of hnRNPC in the response to chemotherapy. We focused our analysis on stage 2 patients, since the benefit from adjuvant chemotherapy is most pronounced in this population. Separation of patients based on hnRNPC expression level revealed that patients whose tumors displayed low expression of hnRNPC benefited significantly from chemotherapy in terms of survival (Figure 7D, right panel, p = 0.019), while patients whose tumors had high levels of hnRNPC expression did not show significant benefit (Figure 7D, left panel, p = 0.68). Taken together, the data in Figure 7 identify hnRNPC as a new RBP involved in the response to platinum drug treatment in NSCLC, and suggest that Transite is an effective tool for identifying novel RBPs that contribute to chemoresistance in human cancer patient RNA expression data sets.

## III. Discussion

Despite their crucial role in post-transcriptional regulation of gene expression, the majority of RNA-binding proteins (RBPs) have unknown functions. To help understand the influence of RBPs on their target transcripts, we developed Transite, a computational method for the analysis of the regulatory role of RBPs in various cellular processes for which differential gene expression data, or other relevant gene sets are available. Our analysis is based on the fact that most RBPs recognize short linear oligonucleotide sequences whose overrepresentation can be computed from gene expression data, and that a large collection of pre-existing motif data for RBPs has been compiled in publicly available databases [45,46].

It is important to note that Transite, in its current form, has significant limitations. First, not all RBPs have strong motif preferences that are amenable to this type of motif-based analysis. Furthermore, there may be considerable redundancy in motif recognition by different RBPs, making prediction of a single RBP challenging. Moreover, the *in vitro*-derived motifs for RBPs may not always reflect *in vivo* binding preferences. These caveats have raised questions about the ability of consensus motifs and PWMs to uniquely predict individual RBP mRNA targets *a priori* on a genome-wide scale, and have led to the development of more sophisticated approaches for predicting specific RBP RNA targets [7,47]. In contrast to those approaches, Transite does not attempt to predict specific mRNAs bound by a particular RBP. Instead, Transite simply looks at the statistical distribution of RBP motif representation in sets of expressed genes to infer putative roles for specific RBPs in some biological process, which can then be directly tested experimentally.

By using two approaches to identify non-random distributions of RBP-binding motifs, followed by back-mapping of those motifs onto those of 174 known RBPs, Transite identified 3 RBPs involved in the human DDR which we could further validate based on independent CLIP-Seq data of their known mRNA targets in cells, rather than using motifs derived from *in vitro* sequence libraries. These findings suggest that, although there are limitations to utilizing *in vitro*-derived motifs, Transite serves as a discovery tool for new biology. Moreover, since users can define their own motifs in addition to those from the database, users are able to upload motifs from CLIP-Seq data of their favorite RBP and use that as a means to analyze enrichment in preexisting data sets. As more RBP motifs become available, they will be incorporated in future versions of the Transite analysis platform.

To further demonstrate the utility of Transite, we performed an analysis of human NSCLC patient data and were able to recover previously-known RBP biology and also identify novel sources of RBP-mediated chemoresistance. Well-known players in the DNA damage response such as ELAVL1 and TIA1 were among the top hits in the tumor resistance gene expression data set, showing that our approach is consistent with previous DNA damage response literature. Transite was also able to identify hnRNPC as a new potential modulator of cisplatin sensitivity in NSCLC patients. Experimental validation of the *in silico* prediction further provides independent support for a critical role for hnRNPC in mediating resistance of NSCLC cells to chemotherapy, which was independently correlated with clinical response in an additional NSCLC patient data set.

Transite is a versatile tool that can be used with any type of gene expression data, the only requirements being a list of gene identifiers and some means to separate foreground and background sets or rank the gene list. Examples of the other types of data that are compatible with a Transite style of analysis include: (1) searching for RBP motif enrichment in 5′ or 3′-UTRs of genes whose translational efficiency changes in response to some stimulus as measured by ribosome or polysome profiling; (2) searching for enrichment of RBP motifs in mRNAs that are localized to specific sub-cellular compartments; (3) *de novo* motif analysis in the entire mRNA of gene expression changes upon knockdown of a nuclease of unknown function. The Transite website (https://transite.mit.edu) makes this tool accessible to a broad group of scientists, provides a means by which the large body of pre-existing gene expression data from microarray and RNA sequencing experiments, for example, can be further leveraged to identify changes in mRNA expression associated with specific RBPs, and reveals potential insights into how RBPs may contribute to the concerted regulation and function of specific cellular processes.

## IV. Materials and Methods

### A. Differential gene expression analysis

Differential gene expression analysis for data sets used in this manuscript was performed with the R/Bioconductor package *limma* [48]. A linear model was fit to each row of the log_2_-transformed expression value matrices, where rows correspond to transcripts and columns correspond to samples. An empirical Bayes method was used to obtain the magnitude and significance of the log fold change between sample groups for each transcript [49]. Raw p-values were adjusted using the Benjamini-Hochberg procedure [50].

### B. Motif databases

Transite incorporates sequence motifs of RBP binding sites from two databases: CISBP-RNA, a catalog of inferred sequence preferences of RNA binding proteins [45], and RBPDB, a database of RNA-binding specificities [46]. Together these contribute 174 sequence motifs of varying lengths (between 6 and 18 nucleotides). All motifs were obtained using *in vitro* techniques for determining RNA targets. The majority of motifs were determined by either systematic evolution of ligands by exponential enrichment (SELEX) [11] or RNAcompete [12]. The RNA binding specificities of two further RBPs were obtained by electrophoretic mobility shift assays (EMSA) [51].

### C. Motif representations

Motif descriptions provided from the databases described above were converted from count matrices to position weight matrices (PWMs), obtained by normalizing each nucleotide’s probability at each position by the mean probability of each nucleotide, 25%.

For *k*-mer-based analyses, PWMs were converted to hexamers and heptamers by generating all *k*-mers for which each position has a probability higher than a certain threshold (see Supplementary Methods). In the work presented here, we used a threshold probability of 0.215, which is a stringency level that works well empirically with the motifs from the motif databases.

Laplace smoothing (also known as additive smoothing) is applied to avoid zeros in count matrices before conversion to PWMs. Zeros might occur if the number of sequences on which the position-specific scoring matrix (PSSM) is based, is too small to contain at least one occurrence of each nucleotide per position. In this case, pseudocounts are introduced [52].

### D. CLIP-seq data analysis

The BED files (output from Piranha analysis) for all CLIP-Seq data sets were retrieved from CLIPdb [53]. Read counts were mapped to RefSeq identifiers using a UCSC table with either just 3′-UTR sequences or the entire mature mRNA of all human mRNAs in Hg19 coordinates. RefSeq identifiers were then summarized to gene symbols. For gene symbols with multiple RefSeq identifiers, the one with the maximum counts was taken, as it was assumed this indicated the most highly expressed transcript. This analysis created two gene lists, one where there was binding in the 3′-UTR (3′-UTR targets) or where there was binding in any region of the mRNA (entire mature mRNA targets). These gene lists were then merged with fold change lists from GEO gene expression data set GSE7880. To generate the non-targets list, the entire mature mRNA list was subtracted from the GSE7880 list.

### E. Package and web development

R package development and documentation was streamlined with *devtools* and *roxygen2*, respectively. Core algorithms were implemented in C++. *ggplot2* [54] was used for data visualization.

The website was developed in R with the reactive web application framework *shiny* from RStudio. The components of the graphical user interface were provided by *shiny* and *shinyBS*, which serve as an R wrapper for the components of the Bootstrap front-end web development framework.

### F. Cell culture and colony formation assays

LG1233/T6a cells (mouse lung adenocarcinoma, in the following referred to as T6a) [55] were grown in RPMI-1640 medium supplemented with 10 % fetal bovine serum at 37 °C in a humidified incubator supplied with 5 % CO_2_. Colony formation assays were performed as previously described [27]. Briefly, 48 hours after transfection with siRNAs or pcDNA vectors, cells were treated with either 4 or 8 µM cisplatin or vehicle for 4 hours. Cells were then re-plated in 6-well plates using 1000 mock-treated or 10,000 cisplatin-treated cells per well. In overexpression assays, 500 µg/ml G418 was added to the media to select for cells transfected with pcDNA vectors. After 10 to 14 days, cells were fixed with 4 % formaldehyde and stained with either SYTO 60 (Thermo Fisher Scientific) or modified Wright-stain (Sigma-Aldrich). Colonies were scanned and counted using Odyssey^®^ CLx Imaging System (LI-COR Biosciences).

### G. siRNA transfection

Silencer Select siRNA (Ambion) transfection was performed using Lipofectamine RNAiMAX following manufacturer instructions (Thermo Fisher) with a final concentration of 5 nM. Cells were then treated as described in the previous section.

### H. Overexpression of hnRNPC

pcDNA3.1 vectors expressing FLAG-tagged mouse hnRNPC were generated as follows. First, total RNA was prepared from KP7B (mouse lung carcinoma) cells using RNeasy purification kit (Qiagen) and was used to synthesize cDNAs using SuperScript cDNA Synthesis System (Thermo Fisher). cDNAs were used as templates in PCR reactions using PfuUltra II HF DNA polymerase (Agilent) and the following primers: 5′-GCCCAT**AAGCT-T**ATG<u>GACTACAAAGACGATGACGACAAG</u>GCTAGC-AATGTTACCAACAAGACAGATCCTCGG-3′ (forward) and 5′-GCCCAT**TCTAGA**TTATTAAGAGTCATCCTCC-CCATTGGCGCTGTCTCTG-3′ (reverse). Restriction sites for HindIII (in forward primer) and XbaI (in reverse primer) are in bold. Sequences encoding FLAG are underlined. The PCR products were cleaved with the indicated restriction enzymes (New England BioLabs Inc), purified (QIAquick PCR Purification Kit, Qiagen) and cloned into pcDNA3.1 vectors. The integrity of the plasmids were confirmed by sequencing (Eton Bioscience, Inc.).

### I. Immunoblotting

Cells were harvested 24 (siRNA-transfected) or 48 (pcDNA vectors-transfected) hours after cisplatin treatment and re-plating. Cells were then lysed in RIPA buffer and subjected to standard SDS/PAGE electrophoresis and transferred to nitrocellulose membranes. The membranes were immunoblotted with antibodies against hnRNPC (ab10294, Abcam Inc., Cambridge, UK) and γ-tubulin (Sigma-Aldrich) following manufacturers instructions.

## Availability

The Transite website is available at https://transite.mit.edu. For workflow integration and advanced analysis, the Transite functionality is also offered as an R/Bioconductor package at https://doi.org/10.18129/B9.bioc.transite. The Transite source code is hosted on GitHub (https://github.com/kkrismer/transite).

## Acknowledgements

We wish to thank members of the Yaffe, Hemann, and Burge labs for helpful advice and discussions. Additionally, we thank Anne E. van Vlimmeren for feedback on the manuscript.

## Author contributions

Conceptualization, I.G.C. and M.B.Y.; Methodology, K.K., A.G., I.G.C., and M.B.Y.; Software, K.K., A.G., and T.B.; Validation, S.V., M.A.B., Y.W.K., and E.D.H.; Formal Analysis, K.K., A.G., and D.A.A; Investigation, K.K., S.V., M.A.B., and E.D.H.; Resources, M.B.Y.; Data Curation, K.K.; Writing - Original Draft, K.K.; Writing - Review & Editing, K.K., Y.W.K., E.D.H., T.B., D.A.A., A.H., B.A.J., I.G.C., and M.B.Y.; Visualization, K.K.; Supervision, A.H., B.A.J., I.G.C., and M.B.Y.; Funding Acquisition, K.K., A.G., T.B., I.G.C., and M.B.Y.

