## Supplementary Methods for "Transite: A computational motif-based analysis platform that identifies RNA-binding proteins modulating changes in gene expression"

### 1 Motif representations

Position specific scoring matrices (PSSM) are used to represent sequence motifs. Transite inherits PSSMs describing RBP binding sites from two sources (see section on motif databases in the main text). Motif databases provide PSSMs in one of three types: Position frequency matrices (PFM), position probability matrices (PPM), or position weight matrices (PWM). Internally, Transite algorithms work exclusively with PWMs in order to make subsequent calculations more efficient.

The elements of a PFM represent absolute count of each nucleotide at each position. PPMs and PWMs are derived from PFMs as follows:

$$\overbrace{\begin{pmatrix} & \text{A} & \text{C} & \text{G} & \text{U} \\ 1 & x_{1,1} & x_{1,2} & x_{1,3} & x_{1,4} \\ 2 & x_{2,1} & x_{2,2} & x_{2,3} & x_{2,4} \\ \vdots & \vdots & \vdots & \vdots & \vdots \\ n & x_{n,1} & x_{n,2} & x_{n,3} & x_{n,4} \end{pmatrix}}^{\text{PFM}} \overset{f_1}{\rightsquigarrow} \overbrace{\begin{pmatrix} & \text{A} & \text{C} & \text{G} & \text{U} \\ 1 & y_{1,1} & y_{1,2} & y_{1,3} & y_{1,4} \\ 2 & y_{2,1} & y_{2,2} & y_{2,3} & y_{2,4} \\ \vdots & \vdots & \vdots & \vdots & \vdots \\ n & y_{n,1} & y_{n,2} & y_{n,3} & y_{n,4} \end{pmatrix}}^{\text{PPM}} \overset{f_2}{\rightsquigarrow} \overbrace{\begin{pmatrix} & \text{A} & \text{C} & \text{G} & \text{U} \\ 1 & z_{1,1} & z_{1,2} & z_{1,3} & z_{1,4} \\ 2 & z_{2,1} & z_{2,2} & z_{2,3} & z_{2,4} \\ \vdots & \vdots & \vdots & \vdots & \vdots \\ n & z_{n,1} & z_{n,2} & z_{n,3} & z_{n,4} \end{pmatrix}}^{\text{PWM}} \quad (1)$$

The conversion functions $f_1$ (from PFM to PPM) and $f_2$ (from PPM to PWM) are applied to each element of the matrix. From the PFM, containing counts of each nucleotide $j$ at each sequence position $i$, a row normalizaton converts nucleotide counts at each position to nucleotide probabilities:

$$f_1(\boldsymbol{x}, i, j) = \frac{x_{i,j}}{\sum_k x_{i,k}}, \quad (2)$$

where $\boldsymbol{x}$ is a PFM, and $i$ and $j$ are its indices. PPM elements are converted to PWM elements by taking a log ratio of the actual element-wise probabilities against the probability of the average nucleotide at the same position:

$$f_2(\boldsymbol{y}, i, j) = \log_2 \frac{y_{i,j}}{p_j} \quad (3)$$

where $\boldsymbol{y}$ is a PPM, and $p_j$ is the *a priori* probability of nucleotide $j$. In Transite, nucleotides are assumed to be equiprobable ($\Pr(\text{A}) = \Pr(\text{C}) = \Pr(\text{G}) = \Pr(\text{U}) = 0.25$).

**Laplace smoothing:** Laplace smoothing (also known as additive smoothing) is applied to avoid zeros in PFMs (and thereby zeros in PFMs and negative infinite values in PWMs). Zeros might occur if the number of sequences on which the PSSM is based, is too small to contain at least one occurrence of each nucleotide per position. In this case, pseudocounts are introduced [1]. Specifically, 0.25 was added to all cells (i.e., raw counts) of PFMs which had at least one raw count of zero.

**Scoring algorithm:** A given $k$-mer from a candidate transcript sequence may be scored by comparison to a PWM of length $k$. In order to calculate the score, the corresponding PWM weight for each nucleotide at each position of the candidate $k$-mer is summed.

Under an assumption of position-wise independence of nucleotides in their contribution to RBP-binding fitness, a random $k$-mer will have a score of zero, whereas positive and negative scores will correspond to higher and lower than average fitness, respectively.

### 2 Monte Carlo sampling

Permutation tests are a means to determine the statistical significance of a test statistic with an unknown null distribution. Since no assumptions are made about the underlying distribution of the statistic, permutation tests belong to the group of non-parametric tests. The null distribution of the statistic is obtained empirically by calculating all possible values of the statistic by rearrangement of the labels of the observations (data points). Each unique ordering of the labels is called a permutation, hence the name. Labels are categorical variables that subdivide the set of observations into groups, e.g., *treatment* and *control*. In order to build the complete empirical sampling distribution of the test statistic $T$ based on $n$ labeled observations, $T$ needs to be calculated for $n!$ permutations of the observation labels. The upper tail probability of the actual test statistic, i.e., the test statistic $T$ calculated with the actual observations $x$, here denoted $T(x)$, is given as follows:

$$Pr(T(x)) = \sum_{y:T(y)\geq T(x)} Pr(y), \tag{4}$$

where $y$ are the permuted observations.

Since the number of permutations grows factorially with the number of observations, calculating $T$ for all permutations is infeasible even for small $n$. Therefore, instead of building the complete null distribution, a sample of the distribution is picked randomly to determine an estimate of the probability of $T(x)$. This process is called Monte Carlo sampling. The estimate is determined by the empirical cumulative distribution functions (lower-, upper- and two-tailed probability):

$$\hat{Pr}_L(T(x)) = \frac{\sum_{i=1}^{n} \mathbb{1}\left(T(y_i) \leq T(x)\right) + 1}{n+1} \tag{5}$$

$$\hat{Pr}_U(T(x)) = \frac{\sum_{i=1}^{n} \mathbb{1}\left(T(y_i) \geq T(x)\right) + 1}{n+1} \tag{6}$$

$$\hat{Pr}_T(T(x)) = \frac{\sum_{i=1}^{n} \mathbb{1}\left(|T(y_i)| \geq |T(x)|\right) + 1}{n+1}, \tag{7}$$

where $\mathbb{1}$ is the indicator function and $n$ is the sample size, i.e., the number of performed permutations.

One is added to both the numerator and the denominator to avoid p-values of zero when the actual test statistic is smaller than all of the test statistics of the permuted data [2].

A confidence interval around $\hat{p}$, i.e., $\hat{Pr}(T(x))$, can be calculated based on the cumulative probabilities of the binomial distribution. This interval is referred to as Clopper-Pearson interval [3]. The exact confidence limits $c_l$ and $c_u$ satisfy the following equations:

$$\sum_{i=n_1}^{n} \binom{n}{i} c_l^i (1-c_l)^{n-i} = \alpha/2 \tag{8}$$

$$\sum_{i=0}^{n_1} \binom{n}{i} c_u^i (1-c_u)^{n-i} = \alpha/2, \tag{9}$$

where $n_1$ is the number of cases where $T(y_i) \geq T(x)$ (see equation 6). If $n_1 = 0$, the lower confidence limit is 0, whereas if $n_1 = n$, the upper limit is 1.

**Applications in Transite:** Monte Carlo sampling is used to obtain an estimate of the significance of local consistency scores (see section 6.1 for details) in SPMA spectrum classification. In this case, $n$, the number of local consistency scores that are sampled from the null distribution is 10,000,000. In TSMA, the null distribution of motif enrichment values (in matrix-based TSMA, $n$ = 2,000,000) and $k$-mer enrichment values (in $k$-mer-based TSMA, $n$ = 50,000) is obtained by randomly sampling foreground sets from the background set without replacement and recalculating motif and $k$-mer enrichment values, respectively.

**Early stopping:** In order to significantly reduce the execution time of the Monte Carlo sampling procedure without reducing the number of permutations (where they matter), the tests are implemented with an early stopping mechanism. If the observed test statistic is deemed not significant after a certain number of samples from the null distribution (in Transite, this decision is made after 5000 samples), the null distribution sampling stops.

### 3 Combining enrichment of motif-associated *k*-mers

The overall enrichment of a motif for $k$-mer TSMA is calculated as the geometric mean of the enrichments of associated $k$-mers:

$$\bar{e} = exp\left(\frac{1}{n}\sum_{i=1}^{n} \log(e_i)\right), \tag{10}$$

where $\boldsymbol{e}$ is the vector of enrichment values of motif-associated $k$-mers. The sum of logarithms is used instead of the product to avoid arithmetic underflow.

### 4 Methods for combining p-values

The following section describes methods to combine the significance (p-values) of enrichment values of a set of $k$-mers that are associated with an RNA-binding protein. These methods are used to obtain a single p-value for the overall significance of enriched or depleted RBP-associated $k$-mers.

In general, the methods of this section can be applied to combine the results of independent significance tests. They are commonly used in meta-analysis, where the goal is to systematically assess and integrate findings of a number of studies about a common body of research.

The problem can be specified as follows: Given a vector of $n$ p-values $p_1, ..., p_n$, find $p_c$, the combined p-value of the $n$ significance tests. Most of the methods introduced here combine the p-values in order to obtain a test statistic, which follows a known probability distribution. The general procedure can be stated as:

$$T(h, C) = \sum_{i=1}^{n} h(p_i) * C \tag{11}$$

The function $T$, which returns the test statistic $t$, takes two arguments. $h$ is a function defined on the interval $[0, 1]$ that transforms the individual p-values, and $C$ is a correction term.

#### 4.1 Fisher's method

Fisher's method (1932) [4], also known as the inverse chi-square method is probably the most widely used method for combining p-values. Fisher used the fact that if $p_i$ is uniformly distributed (which p-values are under the null hypothesis), then $-2\log p_i$ follows a chi-square distribution with two degrees of freedom. Therefore, if p-values are transformed as follows,

$$h(p) = -2\log p, \tag{12}$$

and the correction term $C$ is neutral, i.e., equals 1, the following statement can be made about the sampling distribution of the test statistic $T_f$ under the null hypothesis:

$$t_f \overset{H_0}{\sim} \chi^2_{2n}, \tag{13}$$

where $n$ is the number of p-values.

#### 4.2 Stouffer's method

Stouffer's method [5], or the inverse normal method, uses a p-value transformation function $h$ that leads to a test statistic that follows the standard normal distribution by transforming each p-value to its corresponding normal score. The correction term scales the sum of the normal scores by the root of the number of p-values.

$$h(p) = \Phi^{-1}(1 - p) \tag{14}$$

$$C = \frac{1}{\sqrt{n}} \tag{15}$$

$$t_s \overset{H_0}{\sim} N(0, 1), \tag{16}$$

where $\Phi^{-1}$ is the inverse of the cumulative standard normal distribution function.

An extension of Stouffer's method with weighted p-values is called Lipták's method [6].

#### 4.3 Mudholkar and George's method

The logit method by Mudholkar and George [7] uses the following transformation:

$$h(p) = -\ln(p/(1 - p)) \tag{17}$$

When the sum of the transformed p-values is corrected in the following way:

$$C = \sqrt{\frac{3(5n+4)}{\pi^2 n(5n+2)}}, \tag{18}$$

the test statistic $t_m$ is approximately t-distributed:

$$t_m \overset{H_0}{\sim} t_{5n+4} \tag{19}$$

#### 4.4 Edgington's method

Edgington's method [8] is an additive procedure to combine p-values.

$$h(p) = p \tag{20}$$

The sampling distribution of the test statistic $t_e$ under the null hypothesis is given by combinatorics:

$$Pr(t_e) = \sum_{r=0}^{\lfloor t_e \rfloor} (-1)^r \binom{n}{r} \frac{(t_e - r)^n}{n!} \tag{21}$$

#### 4.5 Tippett's method

In Tippett's method [9] the smallest p-value is used as the test statistic $t_t$ and the combined significance is calculated as follows:

$$Pr(t_t) = 1 - (1 - t_t)^n \tag{22}$$

### 5 Methods for adjusting p-values

When multiple statistical tests are performed in order to identify non-random events in a large pool of events, it is imperative to adjust either the p-values themselves or the cutoff significance level $\alpha$, which is the probability of making a type I error (incorrectly rejecting the null hypothesis). Failure to do so leads to alpha error accumulation, i.e., many false positives.

Without accounting for alpha error accumulation in the $k$-mer enrichment step, the enrichment values of for example 204/4096 possible hexamers would be deemed significant between randomly chosen sets of sequences (assuming $\alpha = 0.05$). This is a direct consequence of the number of tests (4096 in this case) and the accepted probability of making a wrong decision per test ($\alpha$).

Transite supports several methods to adjust p-values in order to avoid the multiple testing problem, all of which take a vector of p-values $\boldsymbol{p} \in [0,1]^n$ and return a vector of adjusted p-values $\boldsymbol{q} \in [0,1]^n$. The $i$th smallest or largest p-value is denoted by $p_{(i)}$, depending on whether the method belongs to the step-down (ordered from lowest to highest) or step-up (highest to lowest) group. The methods can be categorized according to the definition of type I error they control.

#### 5.1 Familywise error rate controlling methods

The familywise error rate (FWER) is defined as

$$FWER = Pr(V > 0), \tag{23}$$

where $V$ is the number of false positives in $n$ tests (i.e., "the family").

Methods controlling the FWER guarantee that $FWER \leq \alpha$.

##### 5.1.1 Holm's method

The adjusted p-values [10] obtained by Holm's method are defined as

$$q_{(i)} = \max_{j \leq i} \left( \min\left( (n - j + 1) p_{(j)}, 1 \right) \right), \tag{24}$$

where $p_{(j)}$ is the $j$th lowest p-value and thus characterizing Holm's approach as a step-down method.

##### 5.1.2 Hochberg's method

Hochberg's method is the step-up version of Holm's method ($p_{(i)}$ is highest p-value) and is uniformly more powerful [11].

$$q_{(i)} = \begin{cases} p_{(n)} & \text{for } i = n \\ \min\left(q_{(i+1)}, (n-i+1)p_{(i)}\right) & \text{otherwise} \end{cases} \tag{25}$$

##### 5.1.3 Bonferroni's method

Bonferroni corrected p-values [12] are given by

$$q_i = \min(p_i * n, 1). \tag{26}$$

It is the oldest and most conservative correction.

#### 5.2 False discovery rate controlling methods

The false discovery rate (FDR) is defined as

$$FDR = E\left(\frac{V}{V+S}\right), \tag{27}$$

where $V$ is the number of false positives and $S$ the number of true positives in $n$ tests.

Methods controlling the FDR are less conservative than the ones controlling the FWER.

##### 5.2.1 Benjamini and Hochberg's method

Similar to Hochberg's method for controlling the familywise error rate, this method is defined as a step-up adjustment [13]:

$$q_{(i)} = \begin{cases} p_{(n)} & \text{for } i = n \\ \min\left(q_{(i+1)}, \frac{n}{i}p_{(i)}\right) & \text{otherwise} \end{cases} \tag{28}$$

Compared to the FWER controlling method, the multiplier is less conservative ($\frac{n}{i}$ to $n-i+1$), leading to smaller adjusted p-values. This method can be used if the components (i.e., p-values) of $\boldsymbol{p}$ are independent and uniformly distributed.

##### 5.2.2 Benjamini and Yekutieli's method

If there are dependencies among the p-values or if independency cannot be guaranteed, Benjamini and Yekateuli's method [14] can be used instead:

$$q_{(i)} = \begin{cases} \gamma p_{(n)} & \text{for } i = n \\ \min\left(q_{(i+1)}, \gamma\frac{n}{i}p_{(i)}\right) & \text{otherwise} \end{cases} \tag{29}$$

where $\gamma = \sum_{i=1}^{n} \frac{1}{i}$.

### 6 Classification of spectrum plots

Two methods were developed to identify non-random spectrum plots, a local consistency score that quantifies the changes between neighboring bins, and a method based on polynomial regression models.

#### 6.1 Local consistency score

The local consistency score quantifies the local noise of the gradient in the spectrum by calculating the deviance between the linear interpolation of the scores of two bins separated by one other, and the score of the middle bin, for each position in the spectrum. The lower the score, the more consistent the trend in the spectrum plot. Formally, the local consistency score $x_c$ is defined as

$$x_c = \frac{1}{n}\sum_{i=1}^{n-2}\left|\frac{s_i + s_{i+2}}{2} - s_{i+1}\right|. \tag{30}$$

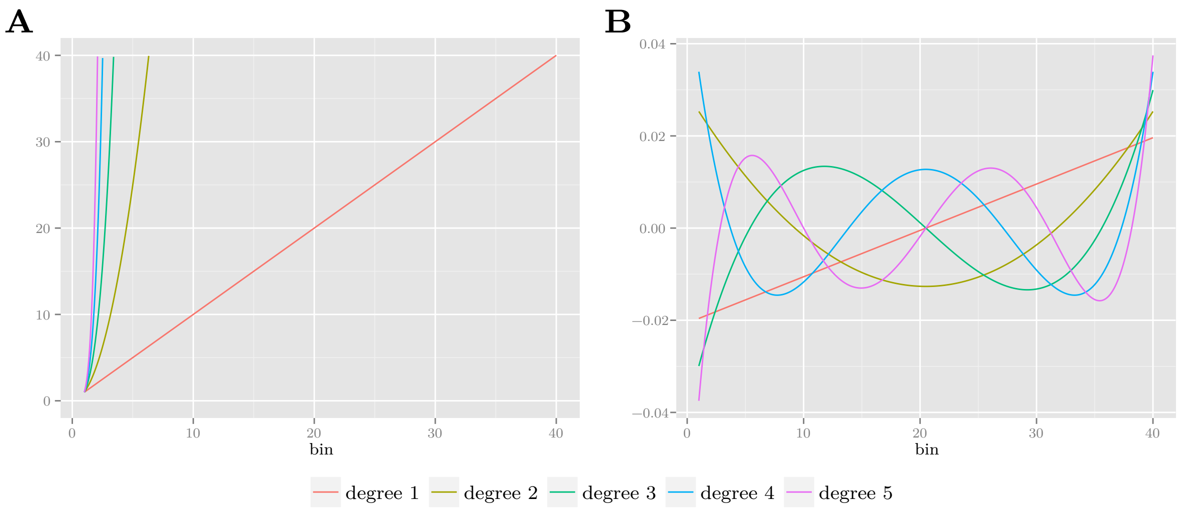


**Ordinary and orthogonal polynomials:** **(A)** The ordinary polynomials of degrees 1 to 5 are highly correlated. Moreover, polynomials of high degree can lead to floating point underflow of model coefficients. **(B)** Correlation between orthogonal polynomials is strongly reduced.

In order to obtain an estimate of the significance of a particular score $x'_c$, Monte Carlo sampling is performed by randomly permuting the coordinates of the scores vector $\boldsymbol{s}$ and recomputing $x_c$. The probability estimate $\hat{p}$ is given by the lower tail version of the cumulative distribution function (see equation 5), where $T$ equals $x_c$ in the equation above.

#### 6.2 Polynomial regression

An alternative approach to assess the consistency of a spectrum plot is via polynomial regression. In a first step, polynomial regression models of various degrees are used to fit $\boldsymbol{s}$, the vector of scores, as a function of $\boldsymbol{b}$, the vector of bin numbers. Then the model that reflects best the true nature of the data is selected by means of the F-test. Finally, the adjusted $R^2$ are calculated to indicate how well the model fits the data. These statistics are used as scores to rank the spectrum plots.

In general, the polynomial regression equation is

$$y_i = \beta_0 + \beta_1 x_i + \beta_2 x_i^2 + \cdots + \beta_m x_i^m + \epsilon_i, \tag{31}$$

where $m$ is the degree of the polynomial (usually $m \leq 5$), and $\epsilon_i$ is the error term. The dependent variable $\boldsymbol{y}$ is the vector of scores $\boldsymbol{s}$ and $\boldsymbol{x}$ to $\boldsymbol{x}^m$ are the orthogonal polynomials of the vector of bin numbers $\boldsymbol{b}$.

Orthogonal polynomials are used in order to reduce the correlation between the different powers of $\boldsymbol{b}$ and therefore avoid multicollinearity in the model. This is important, because correlated predictors lead to unstable coefficients, i.e., the coefficients of a polynomial regression model of degree $m$ can be greatly different from a model of degree $m+1$.

The orthogonal polynomials of vector $\boldsymbol{b}$ are obtained by centering (subtracting the mean), QR decomposition, and subsequent normalization [15].

Given the dependent variable $\boldsymbol{y}$ and the orthogonal polynomials of $\boldsymbol{b}$ $\boldsymbol{x}$ to $\boldsymbol{x}^m$, the model coefficients $\boldsymbol{\beta}$ are chosen in a way to minimize the deviance between the actual and the predicted values. Ordinary least squares is used as the estimation method for the model coefficients. After polynomial models of various degrees have been fitted to the data, the F-test is used to select the model that best fits the data. After a model has been selected, the adjusted $R^2$ is calculated as an additional way to evaluate the goodness of fit.

### 7 Supplementary Figures

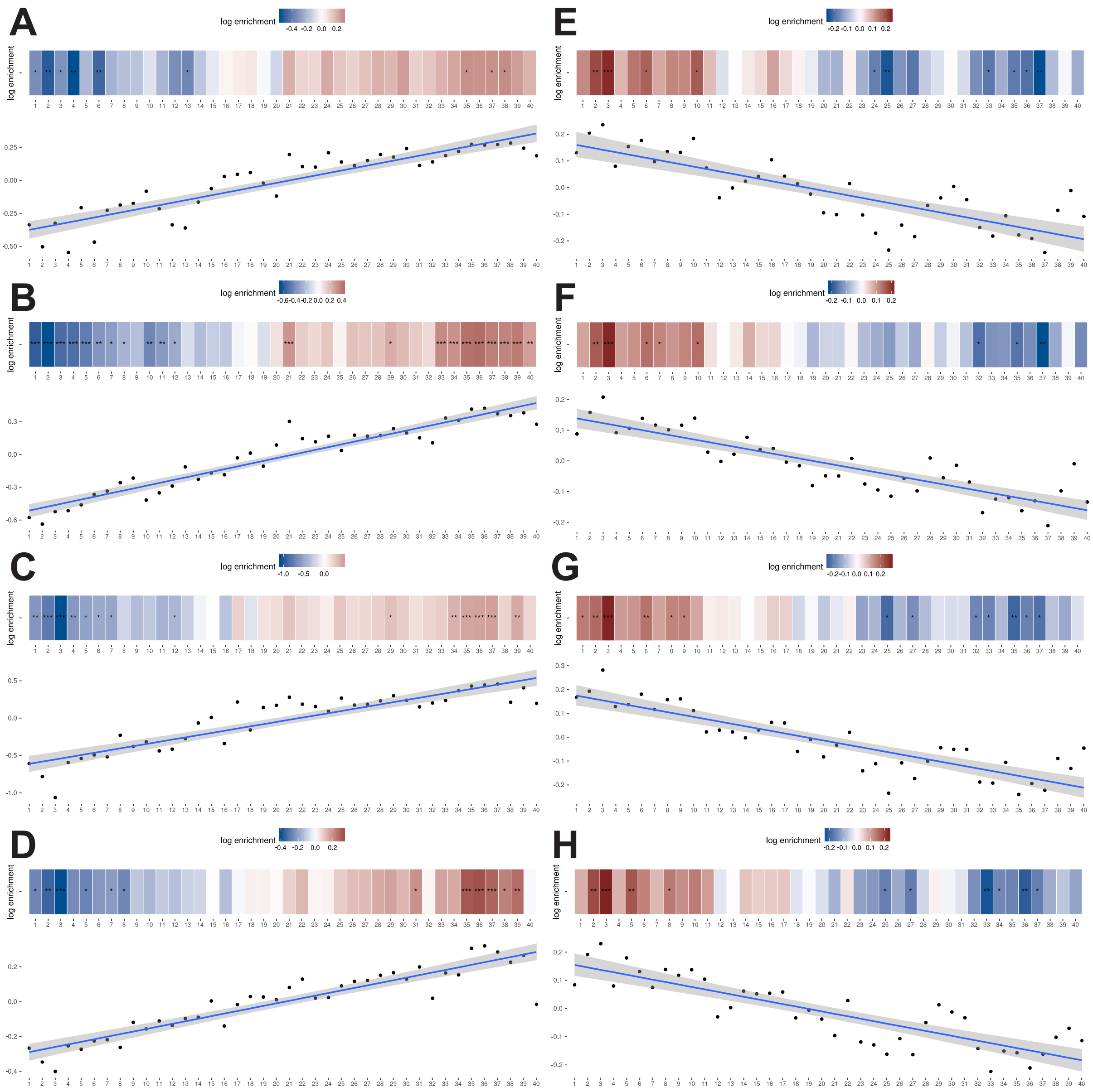


**Supplementary Figure S1: Top spectrum plots of matrix-based SPMA GSE29778.** (**A**) Enriched in upregulated genes: M026_0.6 - hnRNPK (**B**) Enriched in upregulated genes: M043_0.6 - PCBP2/3/4 (**C**) Enriched in upregulated genes: M061_0.6 - SAMD4A, SAMD4B (**D**) Enriched in upregulated genes: M065_0.6 - SRSF9 (**E**) Enriched in downregulated genes: M108_0.6 - ELAVL1, ELAVL3 (**F**) Enriched in downregulated genes: M112_0.6 - ELAVL1, ELAVL3(**G**) Enriched in downregulated genes: M124_0.6 - ELAVL3(**H**) Enriched in downregulated genes: M127_0.6 - ELAVL1, ELAVL3

### 8 Supplementary Tables

| | most enriched hexamers | | most depleted depleted | |
|---|---|---|---|---|
| | **hexamer** | **enrichment value** | **hexamer** | **enrichment value** |
| 1 | CUGGGC | 1.194 | UAAUAU | 0.729 |
| 2 | GGCCUG | 1.191 | AUAUUA | 0.742 |
| 3 | GGGCUG | 1.185 | UAUUAU | 0.749 |
| 4 | GGCCAG | 1.180 | UAUUUA | 0.761 |
| 5 | CCUGGC | 1.179 | UUAUAU | 0.761 |
| 6 | CCCUGG | 1.179 | AAUAUA | 0.767 |
| 7 | CUGGCC | 1.175 | AUUAUU | 0.767 |
| 8 | GCCAGG | 1.175 | UAUAUU | 0.768 |
| 9 | CUGCUG | 1.175 | AAUAUU | 0.770 |
| 10 | GCCCUG | 1.174 | AUAAUA | 0.771 |

**Supplementary Table S1:** Top 10 hexamers statistically significantly ($q < 0.01$) enriched and depleted in upregulated genes after ZFP36 overexpression (GSE53185)

| | most enriched hexamers | | most depleted depleted | |
|---|---|---|---|---|
| | **hexamer** | **enrichment value** | **hexamer** | **enrichment value** |
| 1 | UUAUAU | 1.387 | GGCCCC | 0.458 |
| 2 | UUUAUA | 1.376 | GCCCCC | 0.462 |
| 3 | AUUUUA | 1.375 | GGGGCC | 0.469 |
| 4 | UUAAUU | 1.374 | GGGCCC | 0.480 |
| 5 | UAUAUU | 1.374 | AGGCCC | 0.499 |
| 6 | UUAAAU | 1.372 | GCCCCU | 0.508 |
| 7 | UUUAAA | 1.369 | AGCCCC | 0.510 |
| 8 | AUAUUU | 1.368 | CCCUGC | 0.517 |
| 9 | UUUUAA | 1.366 | CCCCAG | 0.522 |
| 10 | AUUUAA | 1.365 | GGGCCU | 0.523 |

**Supplementary Table S2:** Top 10 hexamers statistically significantly ($q < 0.01$) enriched and depleted in downregulated genes after ZFP36 overexpression (GSE53185)

| | most enriched hexamers | | most depleted depleted | |
|---|---|---|---|---|
| | **hexamer** | **enrichment value** | **hexamer** | **enrichment value** |
| 1 | CACCGG | 1.877 | CAAAUU | 0.698 |
| 2 | CCCCCC | 1.859 | AUUGUG | 0.717 |
| 3 | CGCCCC | 1.730 | UUACUU | 0.737 |
| 4 | CCGGGG | 1.700 | AUGAAA | 0.772 |
| 5 | CAGUAC | 1.643 | UCAUUU | 0.797 |
| 6 | GCCCCC | 1.598 | AAAAAU | 0.802 |
| 7 | CCGCCC | 1.597 | UAAAAA | 0.819 |
| 8 | GGCCCC | 1.591 | AUUUUA | 0.819 |
| 9 | CGGCCC | 1.576 | AUAAAA | 0.821 |
| 10 | CCCUCC | 1.561 | UUAAAA | 0.834 |

**Supplementary Table S3:** Top 10 hexamers statistically significantly ($q < 0.01$) enriched and depleted in upregulated genes after ELAVL1 siRNA knockdown (GSE29778)

| | most enriched hexamers | | most depleted depleted | |
|---|---|---|---|---|
| | **hexamer** | **enrichment value** | **hexamer** | **enrichment value** |
| 1 | AUUUAA | 1.208 | CCCUCG | 0.443 |
| 2 | UAAAAU | 1.197 | CCGGGG | 0.461 |
| 3 | AAAAUA | 1.190 | CGGCCC | 0.468 |
| 4 | UUUAAA | 1.177 | CCCCGG | 0.477 |
| 5 | UUUUAA | 1.158 | UCCCCG | 0.503 |
| 6 | UUUUUA | 1.153 | CCCCGC | 0.504 |
| 7 | | | GCCCGG | 0.506 |
| 8 | | | CGGGGC | 0.515 |
| 9 | | | CGCCCC | 0.518 |
| 10 | | | GCCCCG | 0.519 |

**Supplementary Table S4:** Top 10 hexamers statistically significantly ($q < 0.01$) enriched and depleted in downregulated genes after ELAVL1 siRNA knockdown (GSE29778)

| | most enriched hexamers | | most depleted depleted | |
|---|---|---|---|---|
| | **hexamer** | **enrichment value** | **hexamer** | **enrichment value** |
| 1 | UUUUUU | 1.450 | CCCAAC | 0.582 |
| 2 | GUUUUU | 1.374 | GGGCCC | 0.589 |
| 3 | UUUUUA | 1.364 | GCCCUG | 0.607 |
| 4 | UUUUAA | 1.357 | GCUCCC | 0.617 |
| 5 | UUAAGU | 1.346 | GGCCCA | 0.627 |
| 6 | AAUUUU | 1.327 | GGGGCC | 0.631 |
| 7 | UGUAAU | 1.321 | UGGCCC | 0.633 |
| 8 | UUUUAG | 1.311 | CCCCAG | 0.638 |
| 9 | AAAUUU | 1.310 | CCACCC | 0.639 |
| 10 | UUUUUG | 1.309 | GGCCCU | 0.650 |

**Supplementary Table S5:** Top 10 hexamers statistically significantly ($q < 0.01$) enriched and depleted in upregulated genes in recurrent tumor samples (GSE7880)

| | most enriched hexamers | | most depleted depleted | |
|---|---|---|---|---|
| | **hexamer** | **enrichment value** | **hexamer** | **enrichment value** |
| 1 | CAGGAC | 1.521 | UAGUAA | 0.626 |
| 2 | CCCUGG | 1.513 | UUGAUA | 0.647 |
| 3 | GGACCC | 1.500 | GUUUUA | 0.661 |
| 4 | AGGACC | 1.490 | AAGUUU | 0.720 |
| 5 | CCUGCC | 1.489 | UUUUUU | 0.728 |
| 6 | GACCCA | 1.477 | AUUUGA | 0.729 |
| 7 | AGGGUC | 1.454 | AUUUCA | 0.730 |
| 8 | GGGCCC | 1.435 | AGUUUU | 0.735 |
| 9 | CCCCAC | 1.420 | UAUUUG | 0.746 |
| 10 | GCCCUG | 1.420 | UAAAUU | 0.746 |

**Supplementary Table S6:** Top 10 hexamers statistically significantly ($q < 0.01$) enriched and depleted in downregulated genes in recurrent tumor samples (GSE7880)

### 9 Transite analysis reports

Detailed information on the Transite analysis runs described in the main manuscript are available online.

**HEK293 cells following inducible overexpression of EGFP-ZFP36 fusion protein (GSE53185):**

- $k$-based Transcript Set Motif Analysis report on data set GSE53185
- matrix-based Spectrum Motif Analysis report on data set GSE53185

**HEK293 cells following ELAVL1 siRNA knockdown (GSE29778):**

- matrix-based Transcript Set Motif Analysis report on data set GSE29778
- $k$-mer-based Transcript Set Motif Analysis report on data set GSE29778
- matrix-based Spectrum Motif Analysis report on data set GSE29778

**non-small cell lung cancer patients at diagnosis or at recurrence after cisplatin-based chemotherapy (GSE7880):**

- matrix-based Transcript Set Motif Analysis report on data set GSE7880
- $k$-mer-based Transcript Set Motif Analysis report on data set GSE7880
- matrix-based Spectrum Motif Analysis report on data set GSE7880
- $k$-mer-based Spectrum Motif Analysis report on data set GSE7880
